# Multi-parametric analysis of 58 SYNGAP1 variants reveal impacts on GTPase signaling, localization and protein stability

**DOI:** 10.1101/2020.04.21.053686

**Authors:** Fabian Meili, William J. Wei, Wun-Chey Sin, Iulia Dascalu, Daniel B. Callaghan, Sanja Rogic, Warren M. Meyers, Paul Pavlidis, Kurt Haas

## Abstract

SYNGAP1 is a Ras and Rap GTPase with important roles in regulating excitatory synaptic plasticity. While many *SYNGAP1* missense and nonsense mutations have been associated with intellectual disability, epilepsy, schizophrenia and autism spectrum disorder (ASD), there are many variants of unknown significance (VUS). In this report, we characterize 58 variants in nine assays that examine multiple aspects of SYNGAP1 function. Specifically, we used multiplex phospho-flow cytometry to measure the impact of variants on pERK, pGSK3β and pCREB and high-content imaging to examine their subcellular localization. We find variants ranging from complete loss-of-function (LoF) to wildtype (WT)-like in their ability to regulate pERK and pGSK3β, while all variants retain at least partial ability to regulate pCREB. Interestingly, our assays reveal that a high percentage of variants located within the disordered domain of unknown function that makes up the C-terminal half of SYNGAP1 exhibited LoF, compared to the more well studied catalytic domain. Moreover, we find protein instability to be a major contributor to dysfunction only for two missense variants both located within the catalytic domain. Using high-content imaging, we find variants with nuclear enrichment/exclusion and aberrant nuclear speckle localization. These variants are primarily located within the C2 domain known to mediate membrane lipid interactions. We find that mislocalization is distinct from altered catalytic activity, highlighting multiple independent molecular mechanisms underlying variant dysfunction. Our multidimensional dataset allows clustering of variants based on functional phenotypes and provides high-confidence pathogenicity classification.

## INTRODUCTION

The Ras and Rap-GTPase activating protein (GAP) SYNGAP1 is a 1343 amino acid (AA) protein that contains a core GAP domain and an auxiliary C2 domain essential for its regulation of secondary GTPase targets including Rheb, Rab and Rac^1–8^. As a GAP, SYNGAP1 promotes the dephosphorylation of GTP to GDP by GTPases, thereby inhibiting GTPase signaling by reducing the abundance of their active, GTP-bound form.

SYNGAP1 is one of the most abundant proteins at the post-synaptic density (PSD) complex of excitatory glutamatergic synapses^6–8^. As such, it is well poised to regulate activity-dependent cytoskeletal reconfigurations and AMPA receptor (AMPAR) trafficking associated with both long-term potentiation (LTP) and long-term depression (LTD), processes mediated by Ras and Rap respectively^9–12^. SYNGAP1 binds to the scaffolding proteins PSD95, MUPP1, and SAP102/DLG3, and closely associates with NMDA receptors (NMDARs). Synaptic innervation triggering calcium influx activates Ca^2+^/calmodulin-dependent protein kinase II (CaMK2), which then translocates to the PSD and phosphorylates SYNGAP1, causing its dissociation from PSD scaffolding proteins^2,3,10,13–15^. Its removal causes local increased Ras-activity and AMPAR exocytosis^12,16^. Ras-ERK signaling also promotes expression of immediate early-response genes by activating CREB^17–20^. SYNGAP1 is also implicated in LTD, due to its regulation of Rab5 and Rap1^3,16,21^. In contrast to phosphorylation by CaMK2, phosphorylation of SYNGAP1 by the protein kinases CDK5 and PLK2 shifts its affinity and GAP activity from Ras to Rap, a GTPase that induces AMPAR endocytosis by activating p38^2,11,12,22–24^. SYNGAP1 thus is positioned to act as a regulator of both Ras-ERK-LTP and Rap-p38-LTD signaling.

SYNGAP1 is predominantly expressed in the developing brain and dysfunction of LTD and LTP is believed to contribute to several neurodevelopmental disorders^25–29^. Indeed, reduced function of SYNGAP1 leads to the disruption of several synaptic signaling pathways, and mutations in *SYNGAP1* are associated with intellectual disability (ID), epilepsy, and syndromic SYNGAP encephalopathy^30–32^. *SYNGAP1* (MIM: 603384) mutations have also been identified in individuals with schizophrenia and autism spectrum disorder (ASD)^33–35^. SYNGAP1 dysfunction in model systems replicates disease-associated phenotypes, as a knockdown of zebrafish *syngap1* results in delayed brain development and seizure-like behavior^36,37^. Underscoring the importance of its regulation on LTP/LTD balance, homozygous *Syngap1* null mice die within a week after birth with an increase in neuronal apoptosis, while heterozygous mice have defects in LTP, social-and fear-conditioning and have increased basal activity levels of Rac and ERK^38–43^. Abnormal ERK activation causes epileptic seizures in mice and the Ras-ERK pathway is commonly dysregulated in chronic schizophrenia^44–47^. *Syngap1* heterozygous hippocampus neurons also do not exhibit LTD, presumably through aberrant cofilin activation via Rac^33,48^. Overexpression of SYNGAP1 has the opposite effect and leads to elevated levels of p38, inhibition of ERK, and a reduction of surface AMPARs^49,50^. Additionally, SYNGAP1 has been shown to regulate apoptosis, a process that, like LTD, is regulated by GSK3β^42,51–55^.

While many mutations in *SYNGAP1* are identified as pathogenic early termination/truncating variants, there is a growing number of missense variants of unknown significance (VUS) for which potential contribution to disease development is unclear. To strengthen clinical relevance of *in vitro* findings, and allow for structure-function prediction, functional variomics provides multi-dimensional, deep phenotypic characterization of disease-associated missense variants^56–60^. Here, we use high-throughput multiplex-phospho flow cytometry and high-content screening to measure the impact of 58 variants on protein localization, stability and function in multiple disease-associated signaling pathways, including pERK, pGSK3β, pCREB and pp38 in human embryonic kidney (HEK293) cells. We find diverse impacts of variants and provide deep functional evidence to classify variants as either Likely Pathogenic or Likely Benign.

## MATERIAL AND METHODS

### SYNGAP1 variant selection and cloning

*SYNGAP1* Isoform I (Accession NM_006772.2) was purchased from Genecopoeia (Genecopoeia, EX-H9502). Variants were selected from a variety of sources, including a database of variants isolated in patients without serious pediatric disease (gnomAD^61^), the disease-associated variant database ClinVar^62^, the ASD/ID-associated variant databases SFARI^35^, DDD^63^ and MSSNG, as well as clinical literature sources^30–32,64–68^. We also included a set of likely loss of function (LoF) variants, termed biochemical controls, which are known phosphorylation targets of kinases that regulate SYNGAP1 catalytic ability (CaMK2: Ser1165; CDK5: Ser788, Thr790, Ser817; PLK2: Ser385), as well as variants at sites described as LoF in non-human homologues of SYNGAP1 (Arg485, Asn487, Leu595 and Arg596)^2,22,24,69^. Detailed annotation and sourcing of all variants tested can be found in Table S1. Variants selected were located across the length of the protein, including the four well-annotated domains, including 2xPH, C2, and the Ras/Rap-GAP domain, and within a disordered domain of unknown function (DUF). Variants were generated using three-way Gibson cloning using NEB HiFi DNA Assembly Cloning Kit (NEB, E5520), using a NotI-AscI-digested and purified pENTR backbone (Thermo Scientific, K240020) as well as two PCR-amplified *SYNGAP1* DNA fragments (Start Codon to mutation and mutation to Stop Codon). All variants were then transferred to a custom-made pCAG-mtag-RFP-T-P2A-sfGFP-attr1-ccdb-attr2) destination vector using Gateway cloning (Thermo Scientific, 11791020). Plasmid DNA was isolated using QIAprep Spin Miniprep Kits (Qiagen, 27106).

### Variant assays for stability, function and localization

We investigated whether different missense variants would have impact on protein stability – a major cause of missense variant dysfunction in other genes. To assess this, we used a dual-color RFP-P2A-GFP-*SYNGAP1* construct that would express RFP and GFP at equal rates but as two separate proteins. A reduction in GFP/RFP ratio is indicative of protein instability. To determine variant functional impacts, we assayed the phosphorylation states of several signaling proteins within different, synapse-relevant signaling cascades. We selected assays for pERK1/2 and pCREB as promoters of LTP, and pGSK3β and pp38MAPK as promoters of LTD. Since SYNGAP1 in neurons is highly localized to the PSD complex, we analyzed whether variants exhibited differences in subcellular localization in HEK293 cells. We find that WT SYNGAP1 localized to the nucleus in discrete speckles and used CellProfiler to measure the frequency and shape of speckles, as well as the nucleus/cytoplasm ratio as a metric for nuclear enrichment. Individual variant means, error, N and p-values for each assay are provided in Table S2.

### Cell Culture

HEK293 cells purchased from the American Type Culture Collection (CRL-1573) and were routinely passaged in Dulbecco’s Modified Eagle’s Medium (DMEM) (Millipore Sigma D6046) supplemented with 10% FBS and 100U/mL Penicillin-Streptomycin (referred to as “culture media” hereafter). For all experiments herein, HEK293 cells were used for a maximum of 15 passages. For flow cytometry experiments, cells were seeded at 1×10^5^ per well in 24 well dishes 16-20hrs before transfection with 500ng of expression plasmid using X-tremeGENE 9 at a ratio of 2uL to 1ug DNA. 24h after transfection cells were washed with culture media. 24h later, cells were stimulated for 10 minutes with fresh culture media, then washed once in 1xPBS before treated with Trypsin-EDTA (Gibco, 25200072) for 5 minutes to create a single-cell suspension and then fixed for 10 minutes in 3.2% PFA. Cells were then spun down and resuspended in 100% ice-cold methanol, kept at 4C for 30min before being moved to −20C. For protein localization experiments, cells were seeded at 1.8×10^4^ per well in 96-well black polymer collagen-coated plates (Thermo Scientific) 16-20hrs before transfection. Cells were fixed in 4% PFA with 1:5000 Hoechst-33342 for 20 minutes.

### Antibody Staining and Flow Cytometry

Cells were washed with Flow Cytometry Staining Buffer (FC001, R&D Systems) and then stained in 50ul of Staining Buffer for one hour on ice with the following conjugated antibodies multiplexed at the indicated dilutions: (1) Mouse monoclonal antibody (mAb) anti-Human pS9-GSK-3β-Alexa Fluor 405, 1:50 (R&D Systems, IC25062V), (2) Mousse mAb anti-Human pT202/pY204-pERK1/2-PerCP-eFluor710, 1:200 (Thermo Scientific, 46-9109-41), (3) Rabbit mAb anti-Human pT180/pY182-p38MAPK-PE-Cy7, 1:100 (NEB, 51255), (4) Rabbit mAb anti-Human pS133-CREB-Alexa Fluor 647, 1:50 (NEB, 14001), (5) Mouse mAb anti-Human GAPDH-Dylight680, 1:50 (Thermo Scientific, MA515738D680). Cells were then washed twice with Staining Buffer before being run on an Attune Nxt Flow Cytometer (Invitrogen). Data was recorded using VL-1 (pGSK3β-Alexa Fluor 405), BL-1 (sfGFP), BL-2 (pERK1/2-PerCP-eFluor710), YL-1 (mtagRFP-T), YL-3 (pp38-PE-Cy7), RL-1 (pCREB-Alexa Fluor 647) and RL-2 (GAPDH-Dylight680) channels, which were single-stain compensated. Using FlowJo, Cells were selected using FSC-H/SSC-H and single cells were selected using SSC-H/SSC-A. In-well untransfected control population was selected using BL-1 (sfGFP) and YL-1 (mtagRFP-T) values within spread of values of untransfected control cells, transfected population was selected using BL-1 (sfGFP) values above untransfected to 100-fold above untransfected (Fig S1a).

### High-Content Imaging

Images of 20 fields from each well were collected using the ArrayScan XTI Live High Content Platform (Thermo Scientific). A 20x objective (NA = 0.4, resolution = 0.69 μm) was used to capture widefield images with excitation wavelengths of 386 ± 23, 485 ± 20, and 549 ± 15 nm for imaging of Hoechst, GFP, and RFP respectively. The emission filter wavelengths are 437 ± 25, 520 ± 12, and 606 ± 23 nm respectively. CellProfiler 3.0^70^ was used to analyze the data by identifying nuclei space from Hoechst, cytoplasm space from RFP and nuclear speckles and localization from GFP, respectively. We applied a filter of nuclei and speckle size <1000 pixels as well as cytoplasm size between 200 and 1000 pixels to eliminate imaging artifacts.

### Data Analysis

Relative functional values for each reporter-antibody were obtained by the median of (Individual Transfected Cell Value – Background Value) / (In-well Untransfected Control Median Value – Background Value) values normalized to the sfGFP control = 0 and WT SYNGAP1 = 1 except for stability, where values were normalized to zero as the floor. All data processing, statistical analysis and clustering was performed in Visual Code using python, matplotlib, sklearn and seaborn libraries.

## RESULTS

### ASD/ID-associated missense variants of SYNGAP1 are found throughout the protein

To investigate the impact of variants in multiple domains, we selected 58 variants located throughout the SYNGAP1 protein, including its annotated PH, C2, and GAP, and the C-terminal domain of unknown function (DUF) domain (Fig 1a, Table S1). 28 variants were identified in individuals with ASD/ID and were assigned the primary category ASD/ID. 17 variants were located at sites known to affect SYNGAP1 function and were assigned as biochemical controls (BIOCHEM). 12 variants have not been identified in any patients to date but were found in a database of people without reported pediatric disease (gnomAD^61^). To study the effects of missense mutation on multiple functions of SYNGAP1 we assayed phosphorylated residues of signaling proteins directly downstream of known SYNGAP1 function (Fig 1b). We assayed SYNGAP1 interactions with Ras by assaying phosphorylation states of ERK (pT202/pY204), which is located within the canonical MAPK cascade downstream of Ras^3,43,46,49^ as well as both pS9 on GSK3β and pS133 on CREB, which are known to be indirectly regulated by Ras^47,51,54,71^. To assess SYNGAP1 function towards Rap we assayed pT180/pY182 on p38 MAPK, which is known to be regulated by SYNGAP1 via MUPP1^13,23,49^.

**Figure 1.**
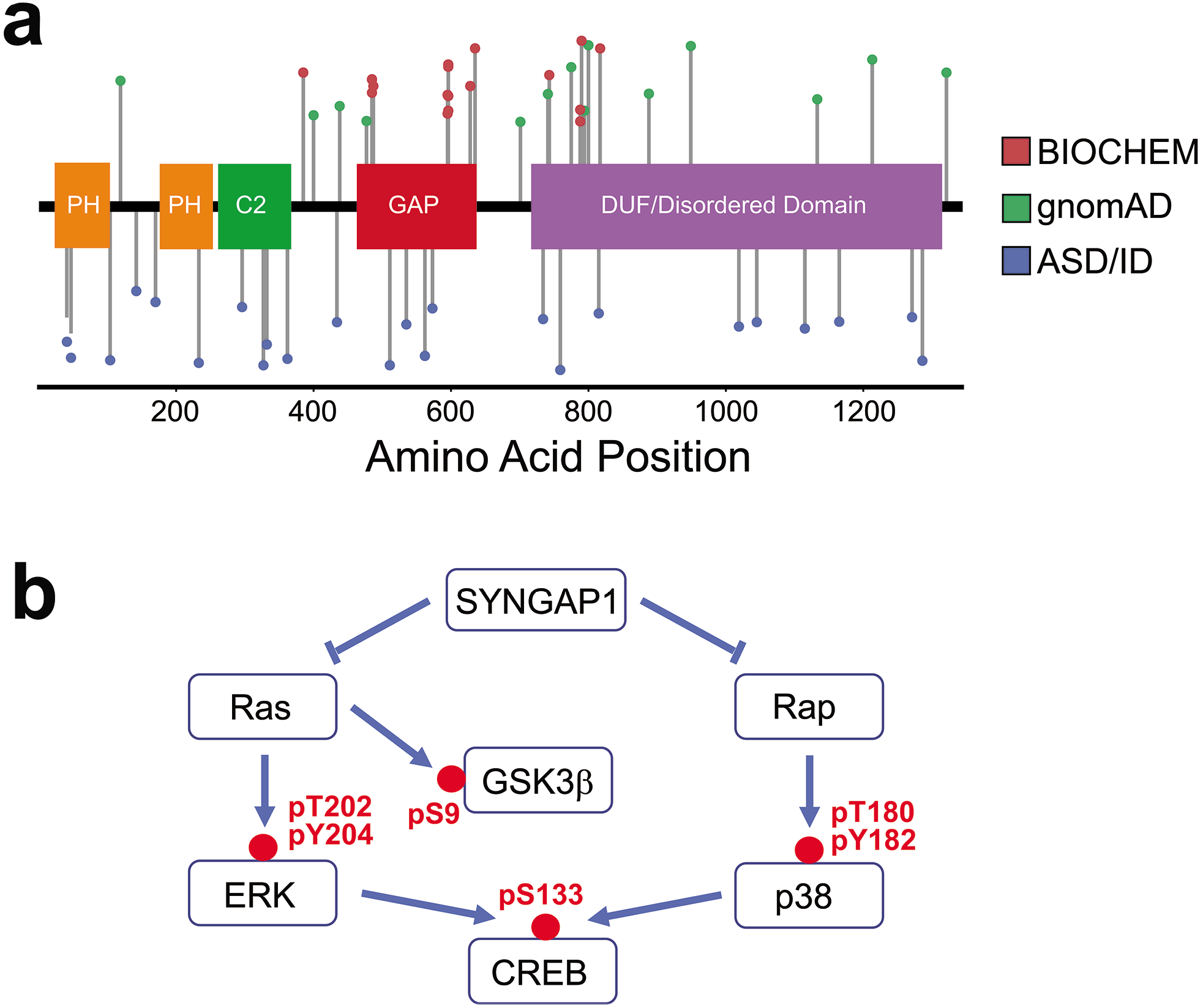
SYNGAP1 variant and functional assay selection. **a.** Distribution of variants assayed in this study across the length of SYNGAP1, including known functional domains. Color indicates whether variants were primarily associated with gnomAD, biochemical control (above functional domains) or ASD/ID (below functional domains). **b**. Simplified SYNGAP1 impact on signaling pathways, including phospho-residues assayed in this study.

### SYNGAP1 missense variants exhibit deficits in inhibiting ERK and GSK3β phosphorylation

We first measured the level of phosphorylated ERK1/2, which correlates with Ras activation and is downregulated by SYNGAP1 (Fig 1b). WT SYNGAP1 reduced phosphorylation of ERK1/2 by 28% compared to levels in untransfected cells. We found variants exhibited a wide range of dysfunction, with 18/58 variants showing complete loss-of-function (LoF), and 22/58 demonstrating partial LoF (Fig 2a). The two missense variants with the lowest (R485A and N487T) and highest (S788A and T790A) functional scores were biochemical controls. R485A and N487T have been previously described as LoF variants, while S788A is a known gain-of-function (GoF) variant and T790A a hypothesized GoF variant based on its proximity to Ser788^2,5,69^. Notably, we observed that across variants, dysfunction was more severe in C-terminal variants, with no variant with less than 50% function being located before amino acid (AA) 485, and all variants past AA800 exhibiting significant LoF (Fig 2b). We find variants located in well-annotated structural domains such as the two PH domains, the C2 domain and the GAP domain to generally have less severe of an impact than variants within the disordered domain of unknown function that makes up the latter half of the protein.

**Figure 2.**
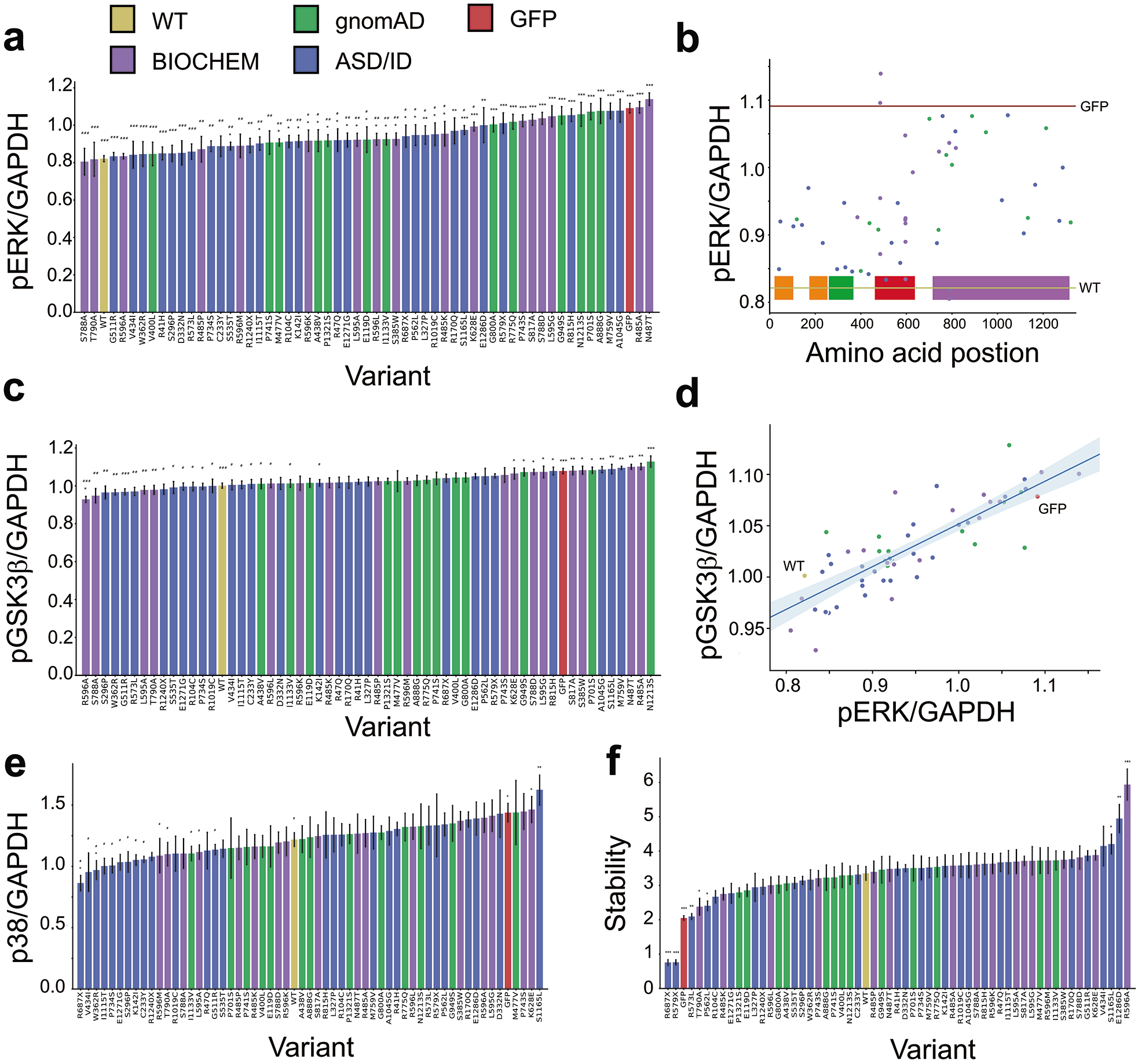
Functional differences of SYNGAP1 variants in ERK, GSK3β, p38 signaling and stability. **a.** Relative pERK/GAPDH value of transfected cells / untransfected cells. Median value of each well averaged across wells, N>=4 wells for all variants. * indicates p<0.05 compared to WT, # indicates p<0.05 compared to GFP by t-test. Color indicates primary variant association. **b.** Scatterplot of pERK vs. pGSK3β functional scores with linear regression. Spearman r=0.81, p=1.5*10^−14^. **c**. Relative pGSK3β/GAPDH value of transfected cells / untransfected cells. N>=4 for all variants. **d**. Distribution of pERK functional scores across the length of SYNGAP1, including domains overlaid at bottom. Red line indicates GFP level, beige line indicates WT SYNGAP1. **e**. Relative pp38/GAPDH value of transfected cells / untransfected cells. Median value of each well averaged across wells, N>=4 wells for all variants. **f**. Protein stability functional value as GFP/RFP ratio of transfected cells. Median value of each well averaged across wells, N>=4 wells for all variants.

Inhibitory effects of SYNGAP1 on the Ras/ERK pathway have been widely described^4,8,38,46,49^. Here, we demonstrate for the first time that SYNGAP1 also regulates a parallel Ras/ GSK3β pathway. Overexpression of WT SYNGAP1 leads to dephosphorylation at Ser9 and thus activation of GSK3β^51^ (Fig 2c). Importantly, the degree of dysfunction of SYNGAP1 variants with respect to ERK inhibition highly correlated with the amount of GSK3β activation, however with lower magnitude (Fig 2d).

P38 MAPK is known to be a downstream target of SYNGAP1 (Fig 1b), and we find that WT SYNGAP1 overexpression significantly dephosphorylates p38 MAPK^13,23^. However, none of the SYNGAP1 variants showed any significant difference compared to WT (Fig 2e), except for LoF of the biochemical control S1165L, which is known to link SYNGAP1 activity to p38 via CAMK2^2^.

### Missense-induced protein instability is not a major cause of dysfunction in SYNGAP1

We next explored whether a measure of protein stability may explain some of our findings, since protein instability is a major mechanism of missense variant dysfunction for other proteins^56,57,72,73^. However, we find SYNGAP1 to be largely resistant to missense-induced instability, with only three of the 58 missense variants tested, R573L, T790A and P562L, exhibiting significant loss of stability, while retaining (Fig 2f). The two early termination variants R579X and R687X were, unsurprisingly, the most unstable, showing ~23% of WT protein abundance. R1240X however, despite missing the last ~100 amino acids of the protein, retained 88% of WT stability. Three missense variants showed significant hyper-stability (S1165L, E1286D and R596A), and in the case of R596A, this increase of ~80% was dramatic.

### Dephosphorylation of CREB independent of ERK/GSK3β dysfunction

The transcription factor CREB is a major regulator of neuronal plasticity and survival and is downstream of several disease-associated signaling pathways, including Ras-ERK^74,75^ (Fig 1b). We find that WT SYNGAP1 significantly decreases levels of pCREB by 26%, similar to its effects on pERK. Moreover, we find that while some variants were fully unable to inhibit ERK, all variants tested suppressed CREB phosphorylation (Fig 3a). 8/58 variants exhibited partial LoF, including known LoF variants at sites phosphorylated by CDK5 (S817A) and CAMK2 (S1165L), while 9/58 variants had a GoF phenotype (again including the known GoF variants S788A and T790A). While functional scores for pCREB were correlated to pERK functional scores (Fig 3b), they were more strongly correlated with the flow-cytometry physical property read-out Side Scatter (SSC), a measure of cell granularity (Fig 3c, d). Changes in SSC are associated with apoptosis and cell death, which is regulated by pCREB levels in neurons^74,76–78^.

**Figure 3.**
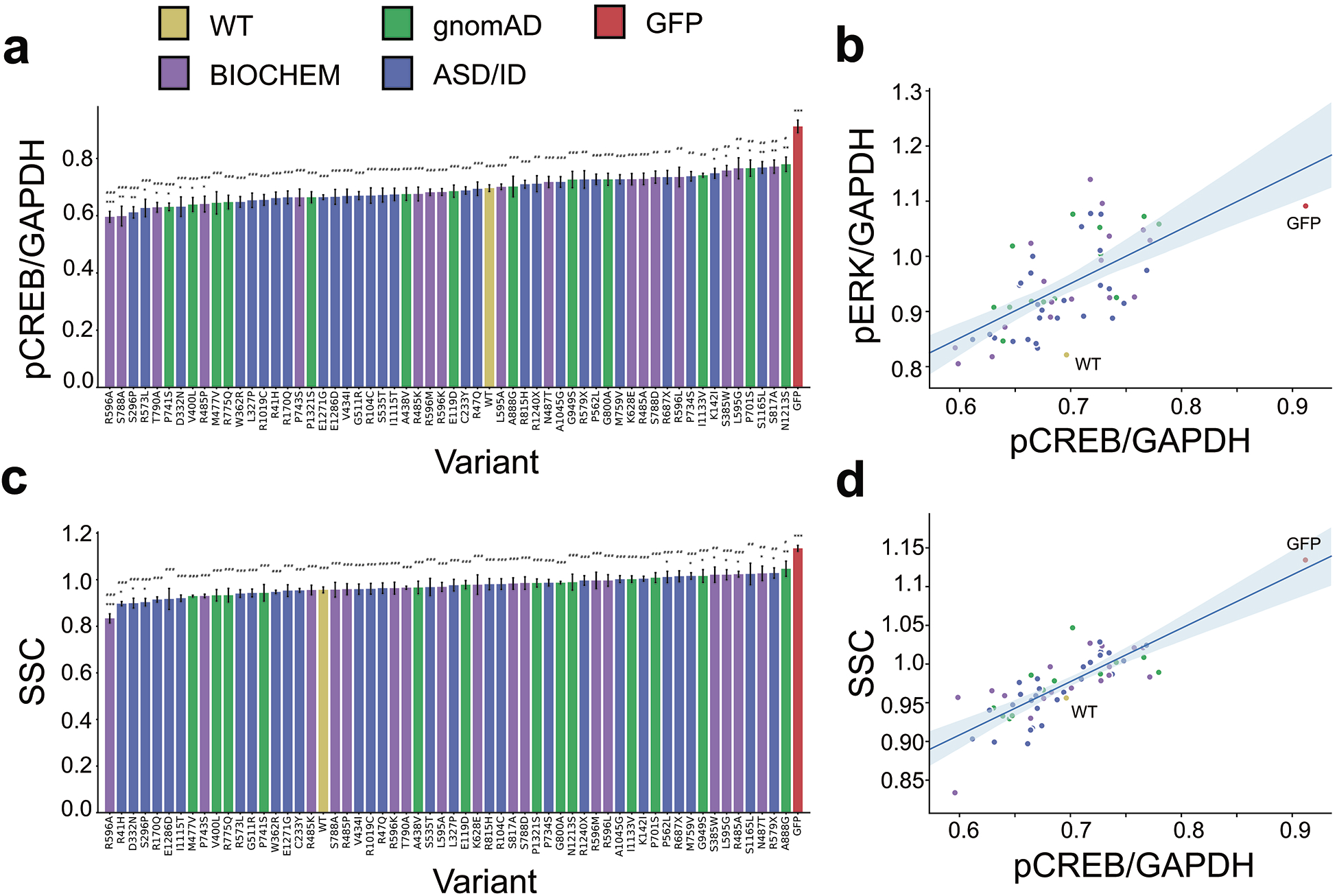
Functional differences of SYNGAP1 variants in CREB signaling and SSC scores. **a.** Relative pCREB/GAPDH value of transfected cells / untransfected cells. Median value of each well averaged across wells, N>=4 wells for all variants. * indicates p<0.05 compared to WT, # indicates p<0.05 compared to GFP by t-test. Color indicates primary variant association. **b.** Scatterplot of pCREB vs. pERK functional scores with linear regression. Spearman r=0.61, p=4.4*10^−7^. **c**. Relative SSC value of transfected cells / untransfected cells. N>=4 for all variants. **d**. Scatterplot of pCREB vs. SSC functional scores with linear regression. Spearman r=0.80, p=5.5*10^−14^.

### Nuclear exclusion is correlated with decreased pCREB

To explore alternate functional pathways that may explain the findings from our pCREB assay, we examined the subcellular localization of overexpressed SYNGAP1 variants. WT SYNGAP1 and variants showed varying levels of either nuclear enrichment or exclusion (Fig 4a, b). 13/58 variants were found to be nuclear enriched, while 29/58 variants showed nuclear exclusion. Notably, the functional scores of variants in our pCREB assay were negatively correlated with their nuclear enrichment, as variants with nuclear enrichment had higher levels of pCREB, while variants with more nuclear exclusion had lower levels of pCREB (Fig 4c). Intriguingly, there was one variant, W362R, which despite strong nuclear enrichment retained the ability to effectively dephosphorylate CREB.

**Figure 4.**
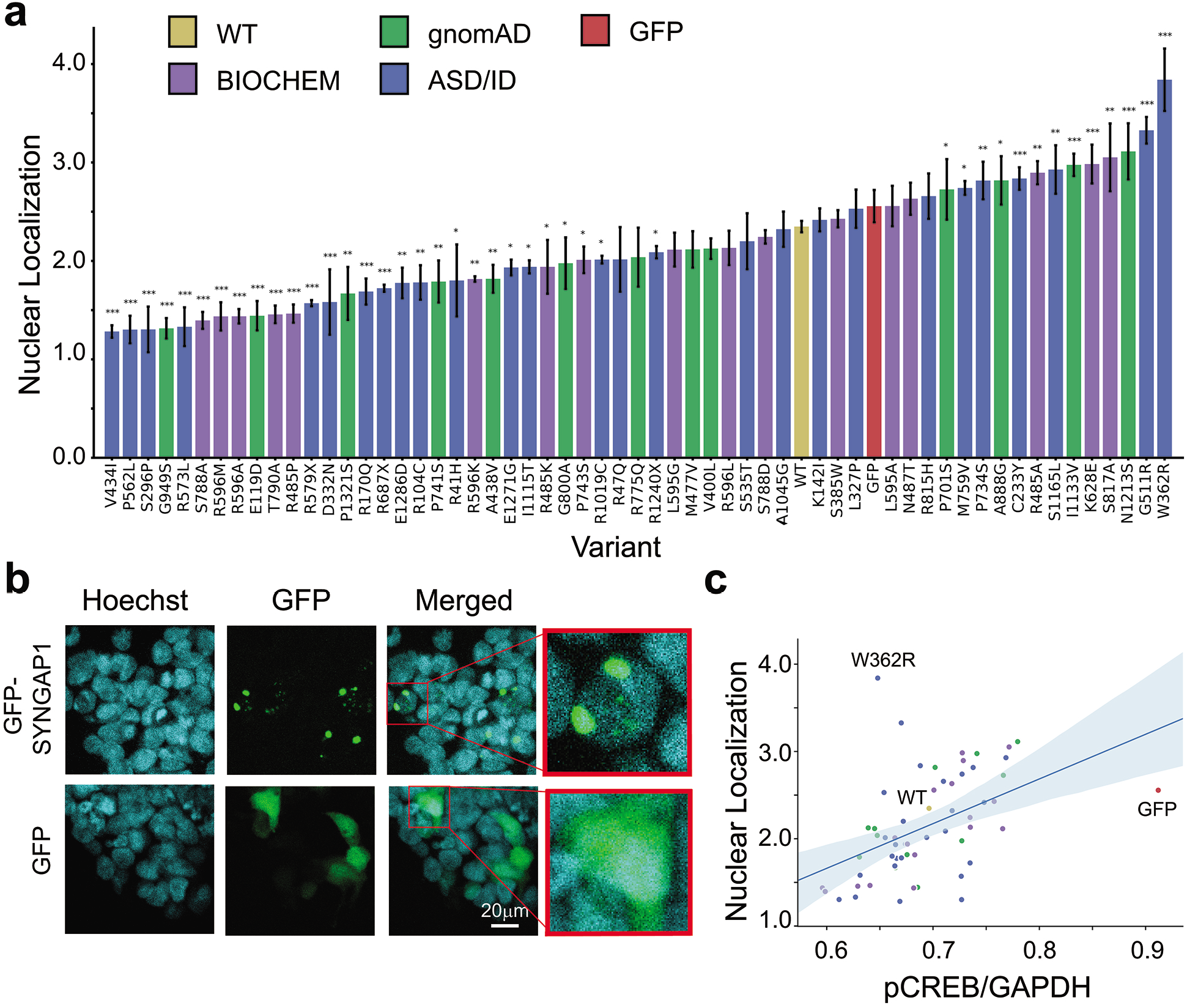
Variants of SYNGAP1 exhibit both nuclear exclusion and enrichment. **a.** Distribution of variant functional scores for nuclear localization measured by sum of nuclear GFP-SYNGAP1 signal over cytoplasmic GFP-SYNGAP1 signal. Mean value of all transfected cells that passed quality filter (see Methods) taken from each well, then averaged across wells. N>=4 wells for all variants. * indicates p<0.05 compared to WT. **b.** Representative images of GFP and GFP-SYNGAP1 localization in HEK293 cells. Cell nuclei were stained with Hoechst (cyan), GFP signal in green. **c.** Scatterplot of pCREB vs. nuclear localization functional scores with linear regression. Spearman r=0.53, p=1.5*10^−5^.

### SYNGAP1 variants exhibited diverse subcellular and nuclear localization phenotypes

We find that SYNGAP1 localizes to puncta (Fig 4b) within the nucleus, and sequence analysis shows that SYNGAP1 contains a poly-histidine repeat from His957 to His966, a repeat that can act as a nuclear speckle targeting domain^79,80^. SYNGAP1 speckled localization has been previously observed^55,81^. We used two parameters that could capture this phenotype – the percent of nuclear puncta that were smaller than 200 pixels (Fig 5a) and the circularity of these speckles, with more irregularly shaped speckles having lower circularity values (Fig 5b). More than half of SYNGAP1 variants showed a decrease in percentage (39/58) and circularity (33/58) of speckles. Six variants, including the seemingly minor substitution E1286D, and a variant lacking the C-terminal domain (R1240X) exhibiting a GoF phenotype of having both a larger percentage of, and in the case of E1286D also more rounded speckles. While speckle percentage and circularity were correlated with each other (Fig 5c), they showed little correlation with other metrics we measured, indicating they might be capturing a distinct function of SYNGAP1. Counter to the functional results in signaling assays, variants with stronger mislocalization phenotypes were localized to the first half of the protein (Fig 5d). L327P, C233Y and W362R, variants close to, or within the C2 domain, exhibited severe LoF for localization, indicating that these domains may be more important in subcellular localization of SYNGAP1 than for its GTPase function.

**Figure 5.**
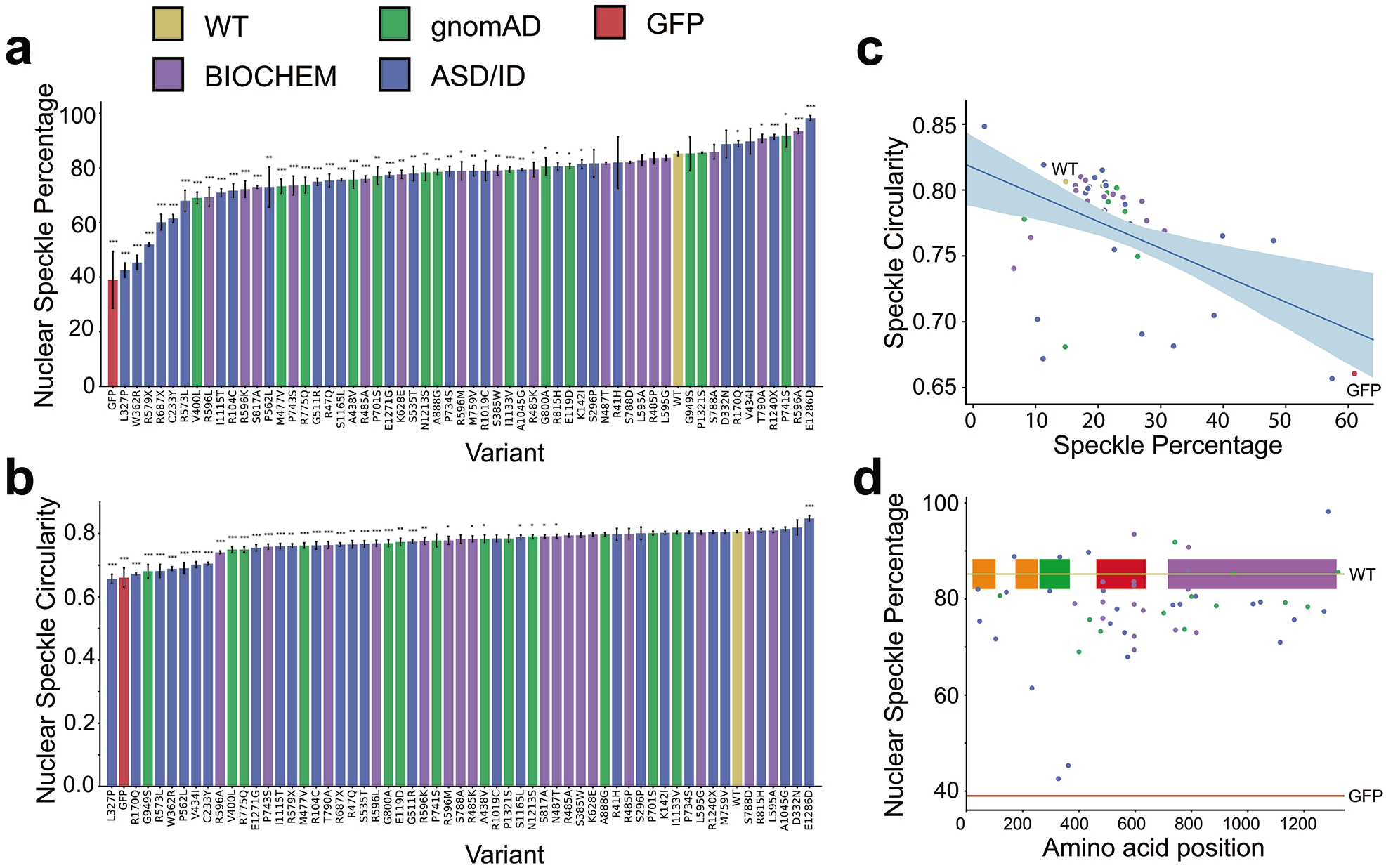
SYNGAP1 localizes to nuclear puncta in HEK293 cells. **a.** Distribution of variant functional scores for nuclear speckle localization measured by the percentage nuclear puncta in transfected cells that passed the nuclear speckle filter and were smaller than 200 pixels (See Methods). Mean across wells with N>= 4 wells per variant. * indicates p<0.05 compared to WT. **b**. Distribution of variant functional scores for speckle circularity with a max value of 1. Mean across wells with N>= 4 wells per variant. **c**. Scatterplot of speckle percentage vs. speckle circularity functional scores with linear regression. Spearman r=0.45, p=4.0*10^−4^. **d**. Distribution of speckle percentage functional scores across the length of SYNGAP1, including domains overlaid at bottom. Red line indicates GFP level, beige line indicates WT SYNGAP1.

### Clustering analyses

Normalized functional scores for all variants across measures are shown in Fig 6a. Overall, we find strong correlation of functional scores across different signaling assays, but not between signaling and localization assays, with the exception of significant negative correlation between nuclear localization and CREB phosphorylation (Fig 6b, Fig S1b). We applied two approaches towards clustering analysis. First, we clustered variants by functional measure scores generating a cluster heatmap to highlight the most closely correlated variants (Fig 6c). Notably, R596A is not closely correlated to any other variant in the study, likely because of its strong GoF phenotype in stability and speckle circularity. Cluster results reveal that missense variants distributed across SYNGAP1 can have very similar effects on a wide range of phenotypes, such as R485A and M759V, or R47Q and R596K, the two most similar pairs of variants in this study.

**Figure 6.**
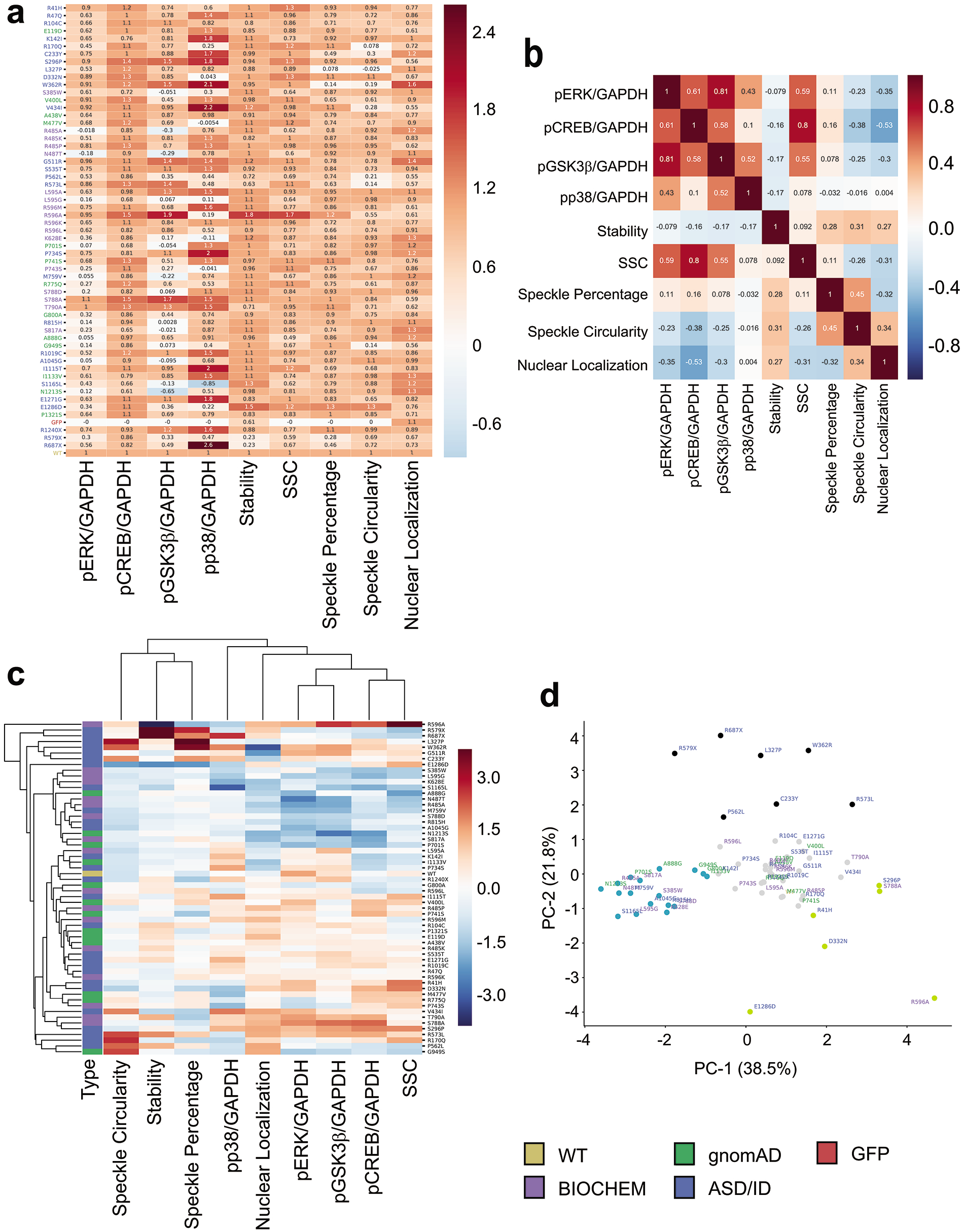
Clustering of SYNGAP1 variants by multi-parameter functional assay scores. **a**. Heatmap of all variant functional scores across assays normalized to WT=1 and GFP=0 except for nuclear localization and stability. Variants are ordered by amino acid position, with early termination variants, GFP and WT at the bottom. **b.** Correlation matrix of Spearman r values. **c**. Hierarchical clustering heatmap of variants and assays calculated from standardized functional scores using Seaborn library. Blue-red gradient indicates standardized variant score in respective assay. **d**. KMeans clustering after PCA performed on all variants in the study. PC-1 accounts for 38.49% of variation in the data, PC-2 accounts for 21.79%. PC-1 primarily accounts for signaling functional scores PC-2 primarily accounts for localization and stability phenotypes (see Fig S1c for PC axis weights).

To illustrate large-scale associations and clusters of variants, we performed Principal Component Analysis (PCA) followed by KMeans clustering on the entire dataset (Fig 6d, Fig S1c). The first PC axis (PC-1) accounts for 38.5% of variation in the dataset and is largely made up phospho-flow scores as well as nuclear localization phenotypes, while the second PC axis (PC-2) features mostly stability and speckle phenotypes, accounting for 21.8% of variation. KMeans clustering reveals four groups of variants: a set of variants that are largely LoF across phenotypes (Turquoise); a second set of variants that are largely WT-like across phenotypes (Black); as well as two additional clusters containing variants with outlier phenotypes in a subset of assays. The two nonsense variants R687X and R579X, as well as the missense variants P562 and R573L are part of the first outlier cluster (Black) as four of the only five variants in this study that exhibited protein instability. This same grey cluster also contains the C2-domain and closely associated variants C233Y, L327P and W362R because of their strong LoF in speckle localization phenotypes. The second outlier cluster (Lime) contains many of the variants that show a GoF phenotype in assays. D332N, R596A, S296P and S788A are GoF in multiple assays, while E1286D is hyper-stable with GoF only in localization phenotypes.

### Multi-parametric pathogenicity prediction

Based on all measures of relative variant function, we devised criteria to predict whether variants were likely pathogenic, likely benign or whether we were unable to make a prediction and a variant remains a variant of unknown significance (VUS). Since no variant exhibited less than 50% function in metrics for pCREB or SSC, we did not use this measure for classification. A variant was classified as Likely Pathogenic (LP) if it had less than 50% or more than 150% function in either 2 out of 3 signaling assays (pERK, pGSK3β or pp38), 2 of 3 localization assays (Speckle Circularity, Speckle Percentage, or Nuclear Localization), or stability. If a variant didn’t fulfill any of these criteria but had less than 50% or more than 150% function in at least 1 assay, we classified it as remaining VUS. If a variant appeared >50% functional in all assays in this study, we classified it as Likely Benign (LB). Based on these rules, we classify 40 variants of SYNGAP1 as either Likely Pathogenic (25/58) or Likely Benign (15/58), while we are unable to make a confident classification determination for 18/58 variants that retained a classification of VUS (Fig 7a). We find that nearly all variants in the C-terminal DUF/Disordered Domain are either Likely Pathogenic (11/21) or VUS (9/21), with only one variant, P741S, being Likely Benign (Fig 7b). We find Likely Pathogenic, Likely Benign and VUS variants across disease association categories, with 12/23 ASD-associated variants being Likely Pathogenic, 5 being Likely Benign and 6 remaining VUS, while we find that of the 13 variants we tested that are present in the non-disease associated reference database gnomAD, 4 are Likely Pathogenic, 4 are Likely Benign, and 5 are VUS (Fig 7c).

**Figure 7.**
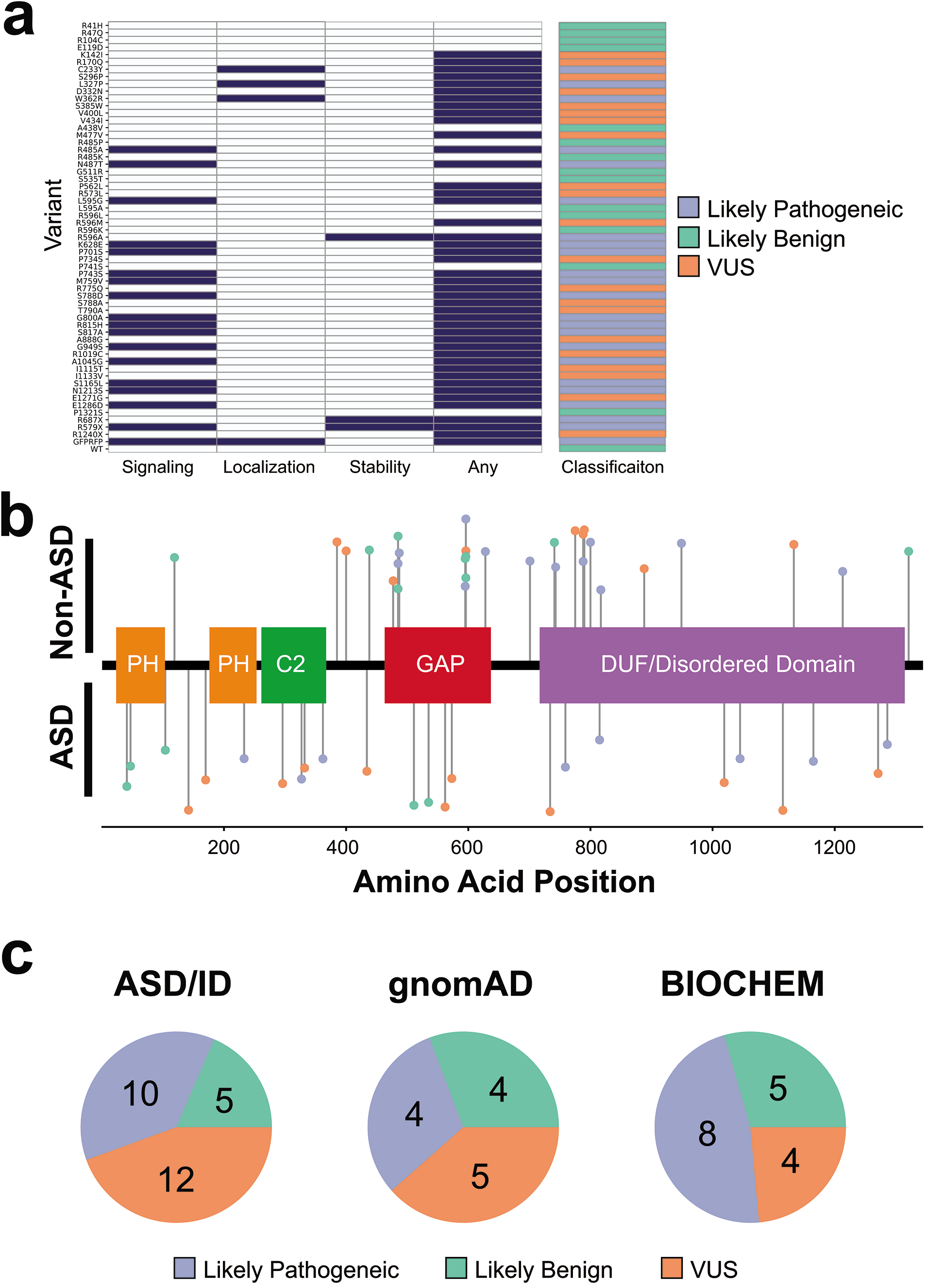
High-confidence pathogenicity prediction of SYNGAP1 variants. **a.** Pathogenicity prediction map for all variants assayed in this study. Blue coloring indicates variant functional score of <50% in at least 2/3 signaling assays, 2/3 localization assays or stability assay. If variants had a functional score <50% in any assay, the “ANY DYSFUNCTION” field is colored blue. Variants that had blue coloring in signaling/localization/stability were classified as Likely Pathogenic (LP), variants that had no blue coloring in any field were classified as Likely Benign (LB) and variants that only had blue coloring in the “ANY DYSFUNCTION” field were classified VUS. **b.** Distribution of pathogenicity prediction across the length of SYNGAP1, including known functional domains. ASD/ID-associated variants plotted below functional domains, non-ASD/ID-associated variants plotted above. **c**. Distribution of pathogenicity prediction across different primary variant association, split by ASD/ID-associated, gnomAD and biochemical controls.

## DISCUSSION

We have characterized the functional effects of 58 variants of SYNGAP1 in nine different functional assays. While the impact on ERK inhibition of biochemical control variants is consistent with previous findings^4,8^, our results show important differences in the effects of mutations in different domains specific to multiple functions of SYNGAP1 and allow us to make high-confidence prediction of variant pathogenicity. We find a range of dysfunction from complete LoF to full WT-level functionality in the impact of SYNGAP1 overexpression (OE) on pERK and pGSK, while all variants assayed retained at least partial function to downregulate CREB. These results indicate that the inhibitory effects of SYNGAP1 OE on CREB are not solely due to its action on Ras/ERK/GSK3β, but through additional pathways leading to a reduction in pCREB. In addition, we find variants affect the ability of SYNGAP1 to localize to nuclear speckles, providing support for a role of this protein in the nucleus.

We find that nuclear excluded variants were better able to dephosphorylate CREB. While CREB is primarily localized in the nucleus both in neurons and HEK293 cells^82^, our results indicate that the initiation of CREB dephosphorylation by SYNGAP1 may take place in the cytoplasm. In neurons, there is evidence that phosphorylated CREB is also localized to axons and dendrites and regulation of CREB phosphorylation plays an important role in neuronal survival, synaptic plasticity, and dendritic growth^83–88^. We identify SYNGAP1 variants deficient in subcellular localization primarily in and adjacent to the C2 domain. In other proteins, C2 domains are known bind to phospholipids at the plasma membrane and missense mutations in the C2 domain have been shown to cause mislocalization^89–93^, matching our results and indicating a similar role of this domain in SYNGAP1. Surprisingly, variants localized within the disordered domain of unknown function (DUF) in the C-terminal half of the protein exhibited the most significant dysfunction in our signaling assays, to a greater extent than variants within the GAP domain. These results support further characterization of the DUF domain and its interaction with SYNGAP1’s GAP domain for understanding missense variant-induced SYNGAP1 dysfunction^94–96^.

We hypothesize that few of the variants in our study contribute to protein instability presumably because as a 1343AA protein, SYNGAP1 is relatively large and can compensate for individual AA substitutions more readily. Indeed, a multi-protein comparative study analyzing missense variants has confirmed the trend that larger proteins tend to be more resistant to missense-induced instability^97^. Two of three instability-inducing variants were located within a span of 11 AA in the center of the GAP domain, and P562L has been previously described as unstable, indicating that only mutations in specific regions of SYNGAP1 may induce instability^32^.

We find two variants that exhibited significant GoF phenotypes across several assays. R596A is localized in highly conserved motif in GAPs, an FLR arginine-finger loop that determines substrate specificity across RasGAPs^69^. Since an alanine substitution of Arg596 will disrupt this motif it is surprising that it would both significantly increase protein stability and retain or enhance function. Further study of the FLR loop in SYNGAP1 and how it differs from other RasGAPs such as p120GAP and NF-1 is warranted. E1286D, on the other hand, is a seemingly minor change from one negatively charged residue to another. While deleterious effects of this conservative substitution have been described in functional roles and thermal stability, it is not well characterized^98,99^. The effect of C-terminal substitutions on SYNGAP1 localization in our assay is supported by the result that removing the last 100 amino acids of SYNGAP1, as in R1240X, produces a GoF phenotype similar to E1286D. The function of the C-terminal domain of SYNGAP1, specifically the α2-isoform used in this study, remains elusive. Despite lacking the C-terminal QTRV-motif found in the α1-isoform, which confers binding to PDZ-motifs, α2 is similarly able to localize to the PSD^15^. Several studies, however, have highlighted that the presence or absence of the QTRV motif is important for SYNGAP-α1 function and its activity-dependent movement from the PSD core^29,100^. Our results indicate that the C-terminal domain of SYNGAP1-α2 also plays a critical role in subcellular localization.

Based on our nine assays, we classify 58 variants, including 16 variants with previous annotations on the disease-associated variant database ClinVar, as Likely Benign, Likely Pathogenic or VUS. Our findings agree with the previous classification of L327P and W362R as Pathogenic, as well as R170Q and S1165L as Likely Pathogenic. However, some of findings disagree with the ClinVar assessment. G511R is present in ClinVar as a single submission with no condition information and has been identified in one individual with epilepsy and autism, while our findings indicate this variant to be likely benign. It is possible that while we used the ClinGEN recommended cutoff of 50% function for classification, milder defects could still be pathogenic, and G511R shows the second-largest nuclear enrichment of all variants in this study (41% increase, p=2.3*10^−8^), providing a possible explanation for the discrepancy. It is also possible that our assays weren’t able to capture certain neuron-specific phenotypes of variant dysfunction or that it is indeed a benign mutation not contributing to patient phenotype. Three variants of SYNGAP1 are present in, and are annotated as Benign in ClinVar: S385W, A1045G and I1115T. While we are unable to make a definitive classification for I1115T, we classify both S385W and A1045G as Likely Pathogenic. S385W is present as a single submission with no condition information on ClinVar, but Ser385 is both a confirmed phosphorylation target for PLK2^22,24^, and S385W has significant LoF in all five of our flow cytometry functional assays. A1045G is one of the most common SYNGAP1 variants in gnomAD and was classified as Benign based on its presence there. However, it shows complete LoF for both pERK and pGSK3β inhibition in our assays, and is also present with >6-fold enrichment in the SCHEMA consortium’s whole-exome database of Schizophrenia-associated variants than in gnomAD, and has been identified in an individual with ASD^34,61,66^. These findings highlight that for low-penetrance variants in multi-genic diseases, heightened presence of an allele in the general population does not necessarily indicate that an allele is Benign. Indeed, large-scale sequencing studies find the presence of seemingly deleterious alleles in otherwise healthy populations at frequencies of more than 100 LoF variants per person^101,102^. Further, other genes in similar multi-genic diseases, such as hypertrophic myocardiopathy, exhibit high-frequency, low-penetrance pathogenic alleles^103^. Here, we are able to add functional evidence and reclassify four variants that were previously annotated on ClinVar only as either VUS or Conflicting (S535T as Likely Benign; P562L, G949S and E1286D as Likely Pathogenic), while for 41 variants we provide the first high-confidence pathogenicity predictions to guide clinicians and researchers.

## SUPPLEMENTAL DATA

Supplemental Data includes one figure and two tables.

## AUTHOR CONTRIBUTIONS

Variant selection and annotation were performed by FM, DBC, SR and PP. Variants were generated by ID and FM. Cell culture and flow cytometry experiments were developed and performed by WMM, WW and FM. High-Content Imaging was performed by WW, WCS and FM. Data analysis was performed by FM. The manuscript was written by FM with editorial assistance from WMM, WCS and KH.

## ACKNOWLEDGMENTS

This work was supported by a grant from the Simons Foundation/SFARI (Grant #573845, Grantees KH&PP) and a CIHR Foundation Award (KH). We would like to acknowledge Manuel Belmadani, Eric Chu and Nathan Holmes for their assistance on variant selection and annotation.

## SUPPLEMENTAL TABLE LEGENDS

**Table S1:** Variants assayed in this study, clinical phenotypes and sources

**Table S2:** Individual results for variants across assays

## SUPPLEMENTAL FIGURE LEGENDS

**Figure S1. Supplemental methods figure for flow cytometry gating and PCA-weights**.

**a.** Gating strategy for flow cytometry data. Cells were selected from SSC-H vs FSC-H, singlets were selected from SSC-H vs SSC-A. Cells with low levels of transfection (up to 20-fold above background fluorescence) were selected from GFP vs. RFP. **b.** p-value matrix of Spearman correlation r values in Fig 6b. **c**. Weights of individual components for PC-1 and PC-2 of PCA analysis performed using sklearn PCA. Larger bars indicate larger contribution of functional scores within the indicated assay for PC-1 and PC-2 axes.

